# Gaze information enhances remote skill transfer of piano performance

**DOI:** 10.64898/2026.05.27.728118

**Authors:** Takanori Oku, Yuto Makimoto, Momoko Shioki, Hideki Koike, Shinichi Furuya

## Abstract

Remote instruction is increasingly used to teach complex sensorimotor skills, yet conventional audio–video communication poorly conveys the fine-grained attentional cues that support expert guidance. This study tested whether real-time bidirectional gaze sharing enhances remote transfer of piano performance skill by restoring joint visual attention between teacher and learner. Twenty-seven conservatory-level pianists were randomly assigned either to a group, in which teacher and learner gaze positions were visualized during online instruction, or to a group receiving otherwise identical instruction without gaze cues. We recorded eye movements with wearable eye trackers and evaluated piano performance using a high-resolution key-motion sensing system. Real-time gaze sharing increased learners’ gaze-pattern similarity to a teacher, which was not evident in the control group. A parallel effect was observed for head-movement similarity. Critically, gaze sharing also reduced variability of the key-descending velocity at the moment of finger–key contact for the right-hand landing after a leap, a feature associated with unstable key-striking velocity. These findings exhibit that gaze information is not merely an auxiliary communication cue but a timing-critical coordination channel for remote motor instruction. By augmenting video-mediated pedagogy with shared attentional dynamics, the proposed system offers a framework for transmitting tacit, high-dexterity skills across distance.

## INTRODUCTION

Transmission of sensorimotor skills beyond constraints of distance is a foundational challenge for continuing education of tacit knowledge such as medical skills and performing arts over spatial boundaries under various situations including pandemic and disaster. To fulfill this requirement, synchronous video communication has become a default medium for teaching complex sensorimotor skills across domains such as sports coaching, surgical telementoring, and music lessons. Yet a persistent barrier remains as to how to preserve the micro-structure of expert guidance when teacher and learner are not co-located. In piano pedagogy, for instance, the gap between basic playing skills and expert-level execution often hinges on nuanced spatiotemporal control of a movement sequence, including anticipatory action preparation and execution of keypress and key-release (Furuya et al. 2011; Winges et al. 2013; Winges and Furuya 2015; van Vugt et al. 2014), together with a teacher’s capacity to assess subtle errors as they unfold. Previous studies described that online instrumental lessons often reduce immediacy and fidelity of teacher–student interaction and make it difficult to evaluate technique and nuanced performance qualities under typical platform constraints (Biasutti et al., 2021; de Bruin, 2021; Daugvilaite, 2021). The literature of online music education also suggests that remote formats can support meaningful learning outcomes, while highlighting heterogeneity in what improves and what remains constrained, especially for performance domains where fine-grained, time-sensitive feedback is central (Lei, 2022). This tension motivates a question in the domain of the human-computer interaction, pedagogy, and sensorimotor learning with direct practical stakes; firstly, which missing informational channels most directly constrain high-fidelity sensorimotor skill transfer online, and secondly, whether interface interventions can restore those channels with measurable benefits.

A compelling candidate is joint visual attention, which represents the ability for a teacher and a learner to share not only what is being attended to, but also when attention shifts and stabilizes around action-relevant events. In co-located instruction, instructors can leverage learners’ gaze and deictic gestures to rapidly establish common ground and to monitor moment-to-moment attentional alignment during joint activity (Richardson et al., 2007; Brennan et al., 2008). In technical skill mentoring, explicitly sharing an expert’s point of regard has been proposed and evaluated as a means to guide trainees’ visual search and improve performance (Leff et al., 2015). In video-mediated instruction, by contrast, directionality cues are often degraded or rendered unreliable because camera gaze does not map onto task-referential gaze, undermining gaze-based grounding and the inference of attention shifts in time (Troje, 2023). Moreover, remote assistance research shows that maintaining mutual awareness of a partner’s viewpoint and focus in a physical workspace is difficult without additional interface cues (Rasmussen et al., 2022). This matters because visual attention in skilled performance is tightly coupled with prediction and action planning rather than reflecting mere interest (Meinz and Hambrick 2010). In pianists, eye–hand coordination metrics including the eye–hand span are systematically modulated by factors such as practice, tempo, and complexity, consistent with anticipatory visual strategies supporting fluent execution (Rosemann et al., 2016). Moreover, the pedagogical importance of micro-structure is not merely conceptual. Our recent study using the high-resolution sensing system shows that subtle movement features (e.g., onset-related noise, key-bottom collision dynamics, and the key-depression acceleration) can carry meaningful performance information that conventional sensing is too coarse to resolve (Kuromiya et al., 2025). Taken together, these lines of evidence suggest that the critical deficit in remote piano coaching may not be “lack of video” per se, but the absence of a high-bandwidth channel linking teacher and learner attention to precisely timed motor events, especially around transitions such as bimanual leaps.

A large body of evidence indicates that sharing gaze can improve coordination when tasks require rapid, time-critical referencing. In collaborative visual search, shared gaze supports highly efficient coordination in time-critical spatial tasks (Brennan et al., 2008). Similarly, in spatial referencing tasks, gaze sharing can be more efficient than verbal communication when rapid consensus is required (Neider et al., 2010). Overlaying expert gaze on instructional materials (i.e. eye-movement modeling examples) can guide attention and improve learning in tasks where selecting relevant information is challenging (van Gog et al., 2009); (Jarodzka et al., 2013). Convergent results appear in sensorimotor domains. In laparoscopic training, gaze-based interventions improve technical skill acquisition and multitasking performance (Wilson et al., 2011), and in sports, training visual attentional control is linked to robust performance under pressure (Vine & Wilson, 2010). Across these literatures, a shared mechanism is that attention guidance can reduce reliance on late corrective control, thereby mitigating high-cost errors that occur at critical moments of action.

Despite these advances, a key issue remains to be unsolved. Previous studies emphasize where attention is directed (spatial reference), whereas failures in high-speed sensorimotor skills are often driven by when attention shifts relative to action (i.e., temporal coordination among gaze, sensory prediction, and timing precision of hand motor control). This distinction is especially salient in bimanual piano tasks requiring rapid leaps in opposite directions, where learners can exhibit a characteristic failure mode. The first keypress after an initial leap tends to be unintentionally “struck” or over-accented, consistent with insufficiently stabilized pre-landing preparation and poorly timed visual anchoring. Here we test whether bidirectional, real-time gaze sharing embedded in a teacher–student lesson mediated by online video communication can (i) increase alignment of learners’ spatiotemporal gaze dynamics with the teacher’s strategy and (ii) yield measurable improvements in keystroke-level microstructure consistent with reduced “over-driving” at the first keypress after a leap, relative to an audio–video-only control. By coupling an interface-based intervention (shared gaze) with hypotheses (temporal attention coupling and anticipatory control) and with fine-grained behavioral outcome measures, our approach aims to strengthen the novel idea that temporal coupling as a target of shared gaze in remote motor instruction, which has significant implication proposing design principles for remote instruction of high-dexterity skills.

## MATERIALS and METHODS

### Participants

In total, 27 pianists who are students at undergraduate or graduate programs of music conservatories (24 female and 3 male, mean age = 23.2 ± 5.5 years, with 20.6 ± 6.5 years of piano training) were recruited and randomly assigned to one of two groups. One group (“**Gaze-Sharing**” group) received instruction with real-time shared gaze visualization, and the other group (“**Control**” group) with no gaze sharing. Fourteen and thirteen pianists were assigned to the two groups, respectively. As the teacher, three professional pianists who won prizes at international piano competitions (mean age = 31.3 ± 7.4 years, 3 females, with 7.7±3.6 years of teaching experience) participated as the teachers in all sessions. Each teacher worked with students in both the Gaze-Sharing and Control conditions (balanced assignment) to control for instructor-specific effects. All participants had normal or corrected-to-normal vision and no reported neurological or motor impairments. Informed consent was obtained from all participants, and the experimental protocol was approved by the local ethics committee of Sony Corporate (Approval No. 24-25-0001).

### Experimental Design

The study employed a between-subjects design with a single experimental factor: the presence of shared gaze visualization during instruction (Gaze-Sharing vs. Control). Students were randomly assigned to either the gaze-sharing condition or the control condition, and each student participated in a single session under their assigned condition (i.e., no crossover between conditions). Each session consisted of a pre-test, training with instruction, and post-test structure. All students first completed a baseline pre-test assessment, then underwent a one-on-one training intervention with a teacher (with or without gaze sharing per group), and finally completed a post-test immediately after training. The Gaze-Sharing group received the real-time gaze visualization during the training phase, whereas the Control group underwent identical piano instruction without any gaze cues. This design allowed us to isolate the effect of real-time shared gaze on learning outcomes by comparing the two groups’ performance improvements from pre-test to post-test. The student-to-teacher pairs were assigned at random, with each teacher participating equally in both sessions. This assignment design ensured that differences between groups could be attributed to the gaze-sharing intervention rather than to characteristics of the participants or instructors.

### Apparatus

We used PupilLabs Neon eye-tracking glasses (Pupil Labs, Berlin, Germany) to capture eye movements from both the student and the teacher in real time. Each participant wore the lightweight eye tracker connected to a companion device (Motorola edge 40 pro: Motorola, Illinois, The United States) for data streaming. The eye tracker captures a gaze position (200 Hz sampling rate), a first-person view video from wearer’s head (30 Hz sampling rate), and head rotation angles in pitch, roll, and yaw direction (90 Hz sampling rate). All data were recorded synchronously by the internal clock of the eye tracker. Gaze data from both teacher and student were transmitted live to a custom interface integrated with the Zoom video conferencing platform (Zoom Video Communications, San Jose, CA). This interface overlaid each participant’s gaze position as a colored cursor on the shared screen with 30 Hz refreshing rate. This enabled to observe one’s gaze positions in real time on the display. The interface is also capable of playing back the teacher’s and student’s synchronized gaze video recording. Each of the gaze video recordings were played back in a vertically stacked arrangement with a slowed-down speed equivalent to 25% of the original speed. Teachers could use this synchronized playback to provide guidance on visual attention related to key striking timings. Prior to each session, the eye trackers were calibrated for each user following the manufacturer’s standard procedure. Each participant was instructed to fixate on the keyboard’s four corners. The experimenter manually corrected discrepancies between actual and estimated gaze positions using the manufacturer’s standard recording application (Neon Companion App).

All lessons were conducted remotely using Zoom. Bidirectional video and audio streams enabled the teacher to observe the student’s playing and communicate instructions. Each student wore the eye-tracker to provide a first-person view of the piano keyboard and the student’s hand movements. The teacher’s video feed (typically showing the teacher’s face and/or a first-person view of their own piano keyboard and hands) was visible to the student. In the Gaze-Sharing condition, the Zoom interface displayed the real-time gaze overlays on top of the video feed; in the Control condition, the Zoom interface was identical except that no gaze cursor was shown. The teacher wore headphones to ensure clear audio and to prevent any audio feedback loops. The Zoom software (version 6.6.10) was used, and all sessions were conducted over stable internet connections to maintain synchronization between audio, video, and gaze data streams (Figure 1. A).

**Figure 1.**
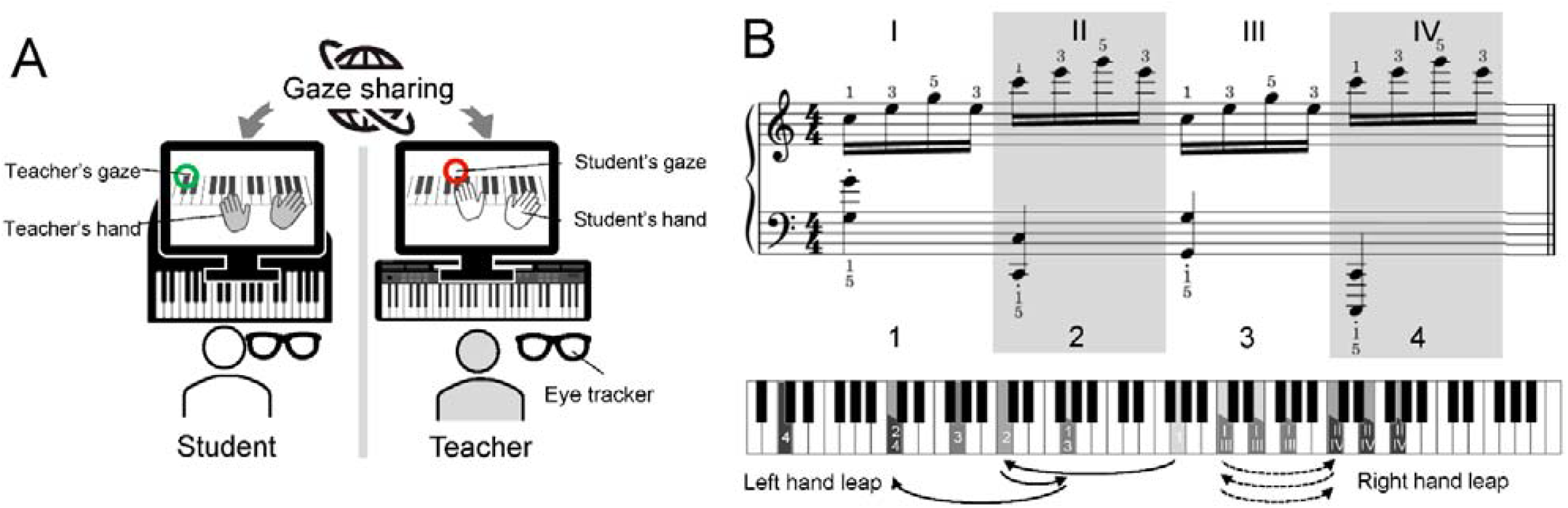
Schematic of the experimental setup. (A) During the instruction session, teacher and student shared gaze information in real time. First-person view videos with gaze positions overlaid as colored cursors were shared between participants. (B) The task was a bimanual piano exercise requiring coordinated hand movements and gaze shifts, in which both hands performed simultaneous leaps in opposite directions.

Students performed on the acoustic piano in the studio environment. An external USB microphone was used to improve audio clarity over the Zoom call. A custom-made contactless piano key motion sensor was equipped on the acoustic piano (Oku and Furuya 2022). The motion data of piano keys were used to synchronize the teacher’s and student’s gaze video recording and evaluate the improvement of the piano performance. The teachers typically did not perform during the session (except for the demonstration performed at the beginning of the lesson), focusing instead on observing and guiding the student. All instructional sessions were recorded (details below) for subsequent analysis.

### Tasks and Procedure

Each participant’s session consisted of a standardized piano task sequence designed to examine the role of gaze sharing in learning. The training task was bimanual piano exercises requiring coordinated hand movements and gaze shifts (Figure 1. B). In particular, the exercise was designed such that both hands had to perform leaps on the keyboard in opposite directions simultaneously, necessitating quick gaze reorientation between the left and right hands. This “contrary motion” design was intended to challenge the student’s ability to allocate visual attention effectively between the two hands.

The session proceeded as follows:

#### Pre-Test (Baseline)

At the start of the session, each student completed a pre-training assessment to establish baseline performance. In this pre-test, the student was instructed to perform a designated bimanual piano task 10 times with 95 bpm tempo. During performance, no guidance or feedback was provided. This task was used to assess the student’s initial spatiotemporal accuracy and natural gaze behavior. The teacher observed silently and took note of the student’s performance and gaze (in the Gaze-Sharing condition the teacher could see the student’s gaze cursor, whereas in Control no gaze cursor was available). No explicit instruction or corrections were given during the pre-test. The pre-test performance provided a baseline for both outcome measures prior to the intervention. *(Duration: approximately 5 minutes*.)

#### Training Session

After the baseline assessment, the student engaged in a focused training session on a specific bimanual piano training task. During this training phase, the student received one-on-one instruction from the teacher. The instruction consisted of ten 3-minute blocks, for a total of 30 minutes. At the beginning of each block, the teacher provided guidance on technique, hand coordination, and strategies to manage visual attention between hands. In the Gaze-Sharing group, the real-time gaze visualization was active: the student could see the teacher’s gaze point on the screen, and the teacher could monitor the student’s gaze point. The teacher utilized this feature by occasionally directing the student’s attention (“Please watch my gaze” or using their own gaze cursor to cue important visual targets) and by observing whether the student was looking at the appropriate hand or section of the keyboard at critical moments. In the Control group, the training was conducted over Zoom in the usual manner without any gaze cues; teachers relied on verbal instructions and demonstrations to guide the student’s focus (e.g., telling the student where to look or what to pay attention to, rather than showing it via gaze cursor). During each training block, students practiced the piece five times, subsequent to the guidance from the teacher. The teacher observed the student’s performance and gaze during practice to determine the next guidance. In order to ensure consistency of instruction across different teachers, a standardized list of instructional content was compiled in advance. In a preliminary experiment involving another six conservatory students, the guidance provided by teachers to students was recorded. The recordings were then reviewed to construct a list of the specific topics to be covered and the appropriate timing for each. Table 1 shows the standardized list of instructional content used in the actual experiment. During the instructional session, the teachers observed the students’ performances and selected the content for each block from the aforementioned list. Both groups thus received the same amount of practice and feedback, with the only difference being the presence or absence of the gaze-sharing overlay. Throughout the training, gaze data and keystroke data were continuously recorded for later analysis, capturing how the student’s visual attention and playing accuracy evolved with practice and instruction.

**Table 1.** The list of guidance contents. The list was compiled in advance to ensure consistency of instruction across different teachers. To construct the list, we recorded the guidance provided by teachers to students in a preliminary experiment. These recordings were reviewed to identify the specific topics to be covered and the appropriate timing for each. During the instructional session, the teachers observed the students’ performances and selected the content for each block from the list.

| No. | Guidance |
| --- | --- |
| 1 | Increase the gaze shift speed. |
| 2 | Look more at the right hand / left hand. |
| 3 | Look at both hands in a balanced way. |
| 4 | Look more at the hands than at the keyboard. |
| 5 | Check the position of the first note, and when starting to play it, check the next position on the left. |
| 6 | Adjust the timing of the gaze shift to the next note position: at the timing of [xx], look at the next left-hand / right-hand position. |
| 7 | During the xx-th leap, your gaze shift speed is too fast / too slow. |
| 8 | During the xx-th leap, look farther ahead. |
| 9 | Orient your head toward the keyboard. |
| 10 | Keep your head as still as possible and look mainly with your eyes. |
| 11 | Reduce head sway. |
| 12 | Increase the speed of head movement. |
| 13 | Move your gaze in an arc-like trajectory. |
| 14 | Move your gaze in a relaxed manner. |
| 15 | Maintain your gaze while paying more attention to the right / left hand that you are not looking at enough. |
| 16 | Maintain your gaze while listening through to the final note. |
| 17 | Maintain your gaze while aligning the key-press and key-release timing of the right and left hands. |
| 18 | Maintain your gaze while making the keystroke of x deeper / shallower. |
| 19 | Try to figure out for yourself why it didn't work out, and then put your plan into action. |
| 20 | Keep it up. |

#### Post-Test

Immediately after the training session, each student proceeded to a post-test to evaluate learning outcomes. Participants performed the trained bimanual piano task 10 times without any real-time instruction or assistance from the teacher. (approximately 5 minutes for the entire post-test.)

Throughout all phases of the session, data was continuously recorded. The Pupil Labs system logged gaze coordinates for teacher and student, and keystroke events (note onsets, etc.) were captured via the instrument or video analysis (described below). This ensured that we collected comprehensive data on both where participants were looking and how accurately they played at each stage of the experiment.

### Data Analyses

We focused on two primary outcome measures to quantify the effects of shared gaze on learning: (1) Gaze Pattern Similarity between teacher and student, and (2) Spatiotemporal Keystroke Accuracy of the student’s performance.

#### Gaze Pattern Similarity

To assess how closely the student’s visual attention aligned with the teacher’s, we computed the cosine similarity between teacher and student gaze patterns. Gaze data from the eye trackers were first processed to generate time-aligned gaze sequences for each participant. Prior to analysis, the gaze time series were resampled (200Hz) and low-pass filtered (20 Hz cutoff frequency). For each performance, we represented time-series horizontal and vertical gaze movements as separate vectors. We then calculated the cosine similarity between the teacher’s and student’s gaze vectors:

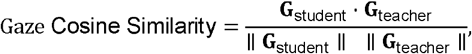

where **G**_student_and **G**_teacher_are the gaze pattern vectors for the student and teacher, respectively. This metric yields values from –1 to 1, where 1 indicates perfect alignment (the student looked at the same places in the same proportions/timing as the teacher), 0 indicates no alignment, and –1 would indicate opposite gaze patterns. An increase in gaze pattern similarity from pre-test to post-test (or a higher similarity in the Gaze-Sharing group compared to Control) would suggest that the student adopted the teacher’s gaze strategy more closely as a result of the training. The time-series vectors for the student’s and teacher’s gazes were extracted from the start of the initial keystroke to the end of the last keystroke in each performance. In the case when the lengths of the gaze vectors of the teacher and student were different, the longer time series was truncated to align with the shorter performance duration. The teacher’s timeseries vectors were derived from the gaze of the first demonstration performance recording.

The similarity of head movements was also computed alongside gaze data to assess how closely the student’s visual attention aligned with that of the teacher. The head rotation time series in each direction were represented as separate vectors and same preprocessing procedures as gaze time series data (resampling, filtering, and time-alignment) were applied to them. We then calculated the cosine similarity between the teacher’s and student’s head rotation vectors.

#### Spatiotemporal Keystroke Accuracy

In sequences involving repeated leaps, keystrokes immediately following a leap (i.e. landing) tend to be unstable. Therefore, we examined performance instability, focusing particularly on the first note following the first leap in the right hand and on the second chord in the left hand (i.e. the first landing of the left hand). We focused on the first leap to avoid a confounding effect of motor learning within the trial. Each student’s performance in the pre-test and post-test tasks was recorded with our custom-made keystroke sensor (Oku and Furuya 2022). Based on the time-varying data of the vertical position of piano keys, we extracted nine spatiotemporal features from piano key displacement data for each keystroke. The parameters were as follows: key velocity at bottom contact (bottom_noise), key velocity at the moment of finger-key contact (onset_noise), key velocity at the moment of keystroke offset (offset_noise), peak striking velocity (peak_vel_des), peak release velocity (peak_vel_asc), peak striking acceleration (peak_acc_des), peak release acceleration (peak_acc_asc), duration of contact between the key and the keybed (pressing_duration), and key velocity at the moment the key passed through the escapement mechanism, which is associated with loudness (vel_escapement) (see Kuromiya et al., 2025 for details). To assess the instability of keystrokes immediately following a leap, we compared the trial-to-trial variability of each keystroke feature between the pre-test and post-test sessions by calculating the standard deviation across trials. We expected that students in the Gaze-Sharing group would show greater reductions in trial-to-trial variability than those in the Control group, if the gaze intervention effectively enhanced their hand coordination and anticipation skills. Trials in which keystroke errors were detected, such as wrong notes, extra notes, or missing notes, were excluded from further analysis.

### Data Collection and Synchronization

All sessions were recorded and synchronized across multiple data streams to facilitate comprehensive analysis. The manufacturer’s standard recording application (Neon Companion App) was used to capture the whole lesson’s video and audio. These recordings included the first-person view video and timeseries of gaze positions from both teachers and students. The video recording (with its audio track) provided a primary reference timeline for the session.

Simultaneously, the Pupil Labs Neon system recorded raw gaze data locally on each participant’s device with high-precision timestamps. These gaze logs (timestamped gaze coordinates for each eye tracker) and timestamps of recording start and end timing were later exported for analysis. Likewise, key movement data collected by the custom-made key movement sensor were used to capture keystrokes and calculate those event timings (with timestamps for onsets and offsets of key striking) (Oku and Furuya 2022). The eye-tracking data and the keystroke data were synchronized using the timestamps. All data streams (gaze, video, audio, keystroke) were thus aligned to a common timeline during data processing, enabling frame-by-frame comparison of what the student played, where they looked, and what feedback was given at any moment in time.

### Post Hoc Feedback and Review

An important feature of our Zoom-based instructional setup was the ability to replay and scrutinize performance footage in real time for educational feedback. We implemented a custom video interface to generate synchronized playback of the teacher’s and student’s gaze recordings. During each session, the student-side interface transmitted the student’s gaze data and keystroke event timestamps to the teacher-side system. The teacher-side interface then integrated the gaze recordings from both participants by aligning them based on the timestamps corresponding to the onset of the first keystroke and the offset of the final keystroke. The resulting synchronized videos were displayed simultaneously in a vertically stacked layout, with the teacher’s and student’s gaze recordings shown in parallel. This configuration enabled direct comparison of temporal alignment and discrepancies in gaze behavior between the different participants.

During the lesson, teachers occasionally utilized Zoom screensharing and the custom-made video interface to review the student’s just-completed performance in detail. For example, after a student finished a difficult segment, the teacher could pause the session and replay the video of that segment (which had been recorded with the gaze overlay in the case of the Gaze-Sharing group) to the student. Using the custom-made video interface, the teacher could pause at a specific frame or play the recording in slow motion, thereby allowing both teacher and student to observe the student’s hand movements and gaze behavior (if available) more closely. This post hoc review was especially useful for highlighting moments where the student’s gaze was suboptimal (e.g., failing to look at the correct hand during a leap) or where a technique lapse occurred too quickly to catch in real time. The teacher would guide the student through the replay, pointing out (verbally and with the aid of the gaze cursor in the Gaze-Sharing condition) what could be improved. Such immediate debriefing via recorded playback was used in both conditions for technical feedback, though in the Control condition only the hand movements and playing could be reviewed, whereas in the Gaze-Sharing condition the visualization of the student’s gaze added an extra layer to the feedback (e.g., “notice how your gaze stayed on your right hand here when your left hand was making the leap; try to shift your gaze earlier next time”). All instances of using slow-motion or pause for feedback were done after the completion of a trial, ensuring that the performance itself was not interrupted by the feedback until a natural break. This approach provided consistency in instruction between groups while leveraging the available data recordings to maximize learning.

In summary, our data collection setup ensured that we had a complete record of each session, with synchronized gaze, performance, and instructional events. This not only facilitated a rigorous post-session analysis of outcome measures but also enriched the instructional interaction through immediate, data-informed feedback. All recorded data were stored on secure servers and were later analyzed using statsmodels (version 0.14.6) library of Python (version 3.12.9) for statistical analysis. Participant identities were coded to maintain confidentiality in all data files.

### Statistics

To evaluate the impact of sharing gaze information, we conducted separate analyses for gaze pattern similarity and spatiotemporal keystroke accuracy. For gaze pattern similarity, linear mixed-effects models (LMMs) were fitted with instructional condition (with vs. without gaze sharing), session (pre-test vs. post-test), and their interaction as fixed effects, and participant-specific random intercepts. The dependent variable was the cosine similarity of gaze patterns between the teacher and the student. The interaction term was used to test whether pre-to post-test changes differed between conditions. When a significant interaction was identified (α = 0.05), post hoc comparisons with correction for multiple comparisons were conducted to assess within-condition changes from pre-to post-test (Benjamini and Hochberg, 1995).

For spatiotemporal keystroke accuracy, linear models were fitted with instructional condition as a fixed effect. The dependent variable was the post–pre difference in the log-transformed trial-to-trial standard deviation of each keystroke feature. The effect of instructional condition was tested to determine whether the magnitude of pre-to post-test change differed between the Gaze-sharing and Control conditions. When a significant condition effect was detected (α = 0.05), follow-up analyses were conducted to test whether the estimated post–pre change in each condition was significant. These tests were corrected for multiple comparisons in the aforementioned manner.

## RESULTS

### Gaze Pattern and Head Movement Similarity

Figure 2 illustrates changes in gaze and head movement similarity relative to the teacher’s performance in the horizontal direction. In the bimanual leap task, eye movements along the keyboard’s longitudinal axis (i.e., the horizontal direction) were predominant. The range of gaze movement in the vertical direction was approximately one-fifth to one-third of that in the horizontal direction. Given that the instructional guidance primarily emphasized the use of lateral eye movements, we therefore focused our analysis on horizontal gaze movements and head movements in the yaw direction. The cosine similarity of gaze pattern between teachers and students was 0.408 ± 0.062 (mean ± standard error) in the Gaze-sharing group and 0.467 ± 0.049 in the control group at pre-test. At post-test, the cosine similarity of gaze pattern between teachers and students was 0.479 ± 0.075 in the Gaze sharing group and 0.378 ± 0.078 in the Control group. The linear mixed-effects model revealed a significant interaction between instructional condition and test timing (β = 0.164, SE = 0.048, z = 3.442, p = 0.001), indicating that changes in cosine similarity differed between instructional conditions. Post hoc comparisons revealed a significant increase in the gaze-sharing condition (p = 0.037), whereas a significant decrease was observed in the control condition (p = 0.004) (Fig. 2A).

**Figure 2.**
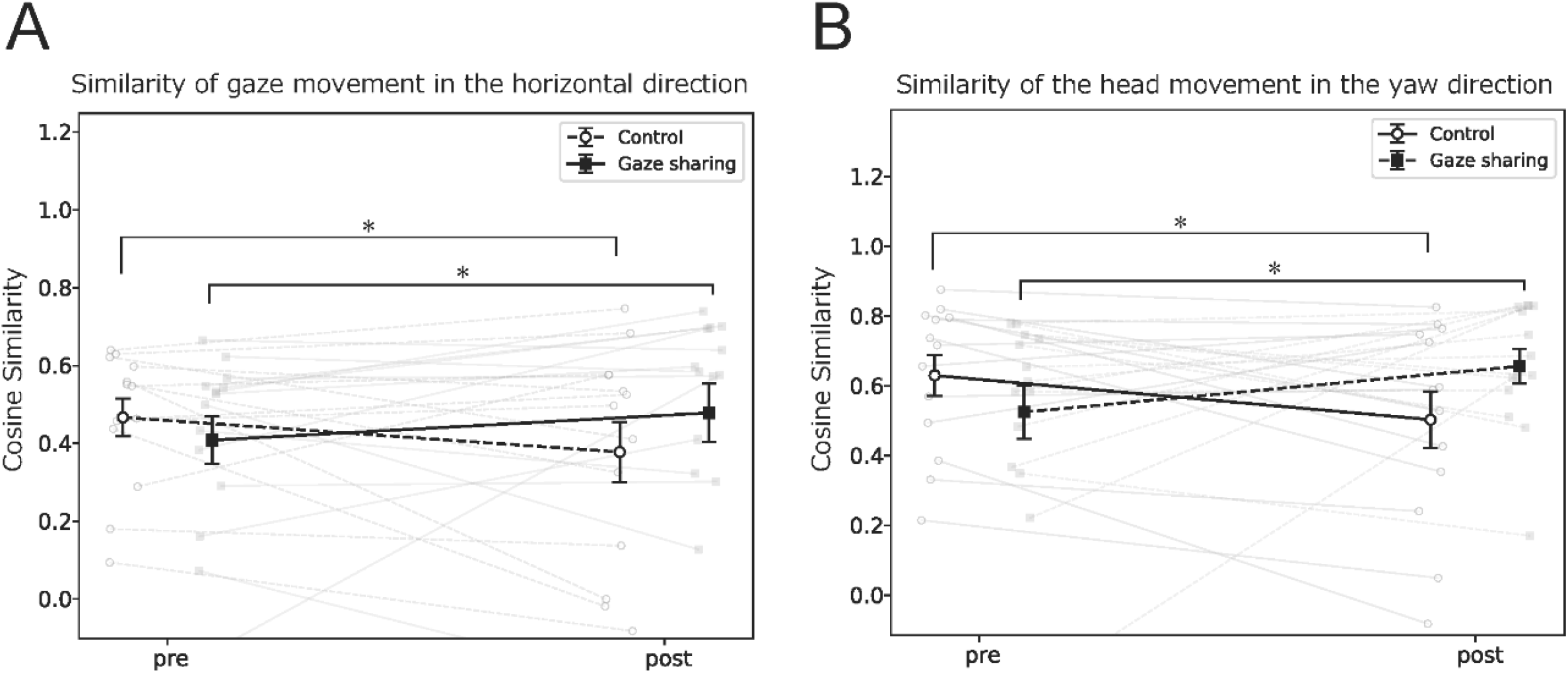
Changes in the gaze movement similarity and the head movement similarity through the intervention. (A) Cosine similarity of the gaze movement between a teacher and a student increased significantly in the Gaze sharing group, whereas it decreased significantly in the Control group. (B) Similarly, cosine similarity of the head movement between a teacher and a student increased significantly in the Gaze sharing group, whereas it decreased significantly in the Control group.

A similar pattern was observed for the head movement similarity. Pre-test values were 0.521 ± 0.030 in the Gaze-sharing group and 0.632 ± 0.026 in the Control group, which changed to 0.656 ± 0.022 and 0.501 ± 0.031, respectively, at post-test. The interaction between instructional condition and test timing was significant (β = 0.264, SE = 0.044, z = 6.048, p < 0.001). Post hoc analysis confirmed a significant increase for the Gaze-sharing condition and a significant decrease for the Control condition (both p < 0.001) (Fig. 2B).

### Spatiotemporal Keystroke Accuracy

Table 2 summarizes the pre–post changes in log-transformed trial-to-trial variability for each keystroke feature following the leap. A significant effect of instructional condition was evident for the trial-to-trial variability of the “onset_noise” at the first note following the first right-hand leap (coef=-0.639, SE = 0.215, z=-2.977, p=0.003). Figure 3 illustrates the corresponding changes in trial-to-trial variability. Post hoc analysis revealed that this variability was significantly decreased from pre-test to post-test in the Gaze-sharing condition (p = 0.001), whereas no significant change was observed in the Control condition (p=0.295).

**Table 2.**
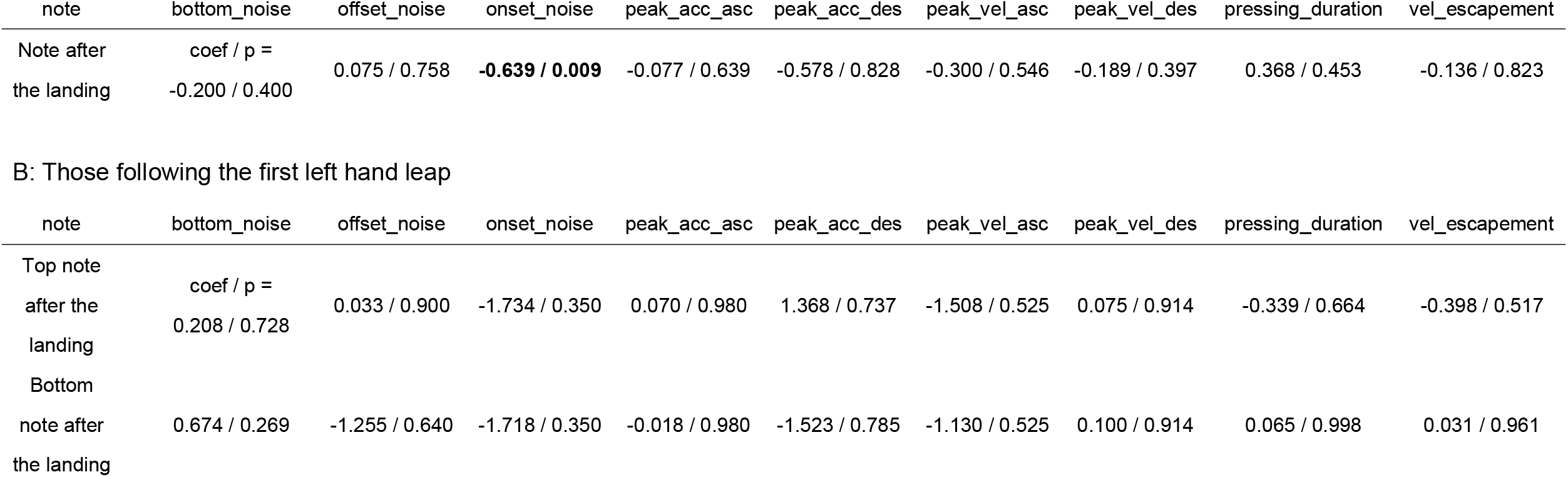
The fixed effect of instructional condition on trial-to-trial variability of spatiotemporal features of keystrokes following the leap. A: Effect of the instructional condition on the trial-to-trial variability (standard deviation across trials) of the spatiotemporal features of the keystrokes following the first right hand leap

| note | bottom_noise | offset_noise | onset_noise | peak_acc_asc | peak_acc_des | peak_vel_asc | peak_vel_des | pressing_duration | vel_escapement |
| --- | --- | --- | --- | --- | --- | --- | --- | --- | --- |
| Note after the landing | coef / p =<br>-0.200 / 0.400 | 0.075 / 0.758 | **-0.639 / 0.009** | -0.077 / 0.639 | -0.578 / 0.828 | -0.300 / 0.546 | -0.189 / 0.397 | 0.368 / 0.453 | -0.136 / 0.823 |

B: Those following the first left hand leap
| note | bottom_noise | offset_noise | onset_noise | peak_acc_asc | peak_acc_des | peak_vel_asc | peak_vel_des | pressing_duration | vel_escapement |
| --- | --- | --- | --- | --- | --- | --- | --- | --- | --- |
| Top note after the landing | coef / p =<br>0.208 / 0.728 | 0.033 / 0.900 | -1.734 / 0.350 | 0.070 / 0.980 | 1.368 / 0.737 | -1.508 / 0.525 | 0.075 / 0.914 | -0.339 / 0.664 | -0.398 / 0.517 |
| Bottom note after the landing | 0.674 / 0.269 | -1.255 / 0.640 | -1.718 / 0.350 | -0.018 / 0.980 | -1.523 / 0.785 | -1.130 / 0.525 | 0.100 / 0.914 | 0.065 / 0.998 | 0.031 / 0.961 |

**Figure 3.**
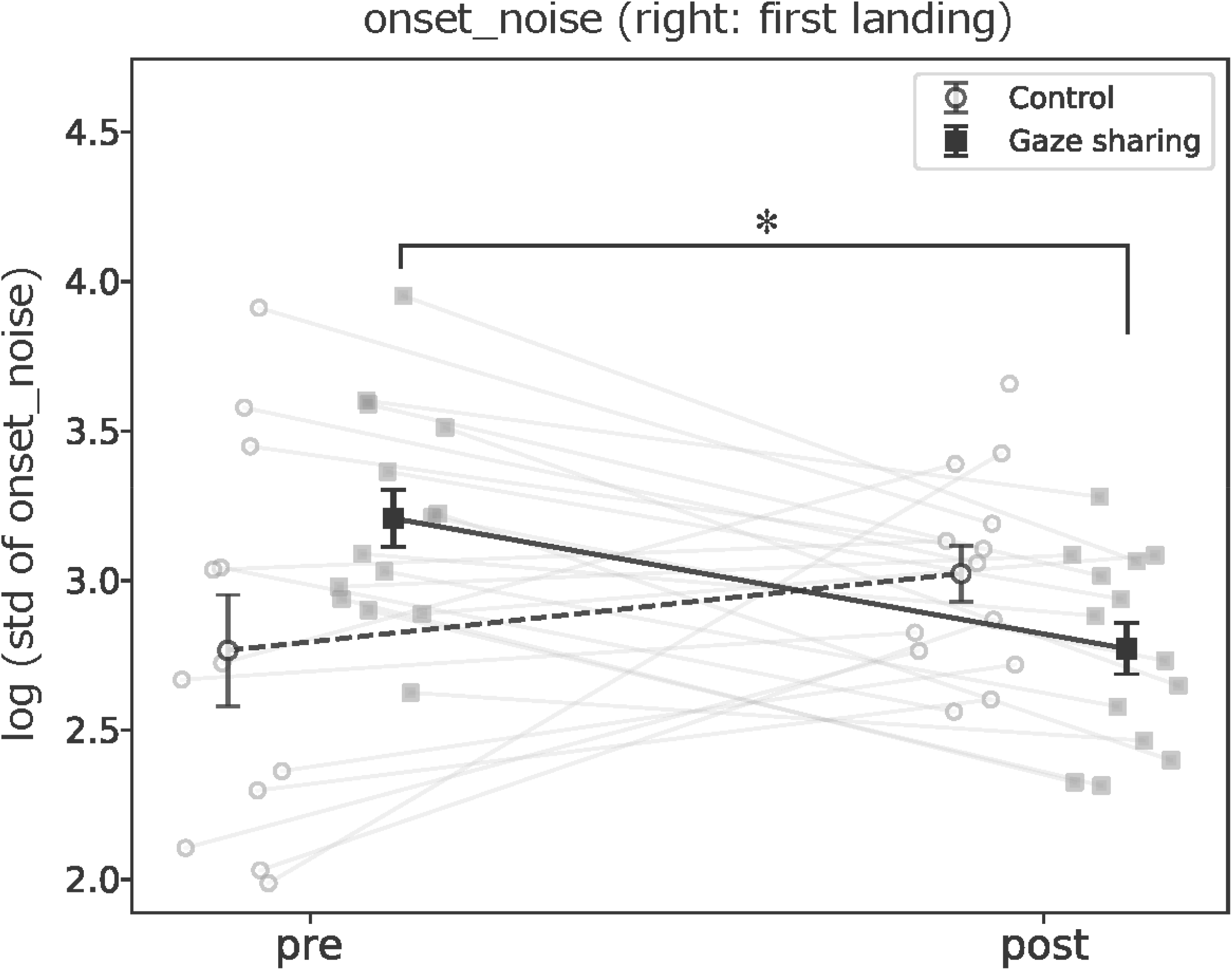
Changes in the trial-to-trial variability of the keypress velocity at the moment of finger-key contact following the leap (onset_noise). There was a significant decrease in the standard deviation of the onset-noise across trials specifically in the Gaze-sharing group.

## DISCUSSION

This study provides evidence that bidirectional gaze sharing can enhance remote transfer of a bimanual motor skill of dexterous piano playing beyond audio–video communication alone. In the gaze-sharing condition, spatiotemporal gaze patterns of the students became more aligned with the teacher’s one through the intervention, whereas the control condition did not show the same directional change. Moreover, the analyses of the keystroke motions support that the intervention is consistent with mitigation of a characteristic motor error in bimanual leap execution, which represents unintended over-accenting at the first keypress after a leap. This was reflected in reduced trial-by-trial variability of the key-descending velocity at the moment of the finger-key collision (i.e. onset noise). This combination of the teacher–learner attentional alignment and the improved nuanced features of the keystrokes provides direct evidence that sharing gaze information in a remote education situation allows for transferring teachers’ skills to a learner, emphasizing the importance of human augmentation technologies for communicating tacit knowledge beyond constraints of distance.

From the perspective of cognitive science and human-computer interaction, our findings sharpen the role of gaze sharing from a generic communication-aid to a timing-critical coordination channel for sensorimotor teaching. In remote collaboration and joint visual tasks, shared gaze can improve coordination by externalizing a partner’s attentional reference (Brennan et al., 2008); (Neider et al., 2010). At the instructional level, previous studies show that visualizing expert gaze can guide attention and improve learning from demonstrations (van Gog et al., 2009); (Jarodzka et al., 2013). Together, the literatures support a coherent prediction for remote piano coaching, which is that restoring joint visual attention, including its temporal structure, should reduce the burden on explicit verbal specification and support rapid diagnosis/correction of attentional timing errors around movement transitions.

The results also resonate with convergent evidence from other high-stakes sensorimotor domains, which exhibits that modifying visual attentional control can produce robust skill gains. In laparoscopic training, gaze-training interventions improve technical skill acquisition and multitasking performance (Wilson et al., 2011). In sports, quiet-eye training improves learning and can protect performance under pressure (Vine & Wilson, 2010). The present work extends these insights to live remote mentorship, where the core deficit is not simply lack of visual information, but the loss of a high-bandwidth channel linking teacher and learner attention to precisely timed motor events. This extension is consistent with emerging work in which experts share gaze via mixed reality to guide novices with reduced verbal effort and improved task performance (Acar et al. 2023).

In conclusion, the present study provided potentials of sharing gaze information between teacher and learner as effective means for enhancing skill transfer in remote skill education of the piano performance. However, several limitations constrain inference and motivate follow-up experiments. First, although the present study quantifies gaze and head movement and links them to the nuanced keystroke movements, it lacks direct measurement of the whole-body and upper-extremity kinematics that likely contribute to preparation for the leap motion and the emergence of over-accented keypress. Second, additional controls are needed to further isolate which component of the gaze intervention drives learning, such as bidirectional versus unidirectional (from expert to novice) gaze visualization, and real-time overlays versus post-hoc replay only. Third, the present study emphasizes immediate post-training effects, and therefore, retention and far transfer across repertoire and expertise levels should be tested. Finally, to uncover underlying mechanism, future studies should combine kinematics with targeted perturbations to adjudicate whether gaze sharing primarily improves anticipatory planning, error detection, or teacher–student grounding efficiency.

## Acknowledgements

This study was supported by JST CRONOS (JPMJCS24N8) and JST Moonshot R&D (Grant Number JPMJMS2012).

